# Mechanical Causation of Biological Structure: Productive Pulls Produce Persistent Filaments in a Human Fibroblast Model of Matrix Development

**DOI:** 10.1101/2021.09.10.459665

**Authors:** Alexandra A. Silverman, Jason D. Olszewski, Seyed Mohammad Siadat, Jeffrey W. Ruberti

**Author notes:** Jeffrey W. Ruberti, Ph.D. **Email:**. **Author Contributions:** Designed experimental approach: AAS, JDO, SMS, JWR; Conducted Experiments: AAS; Collected data: AAS, JDO; Analyzed data: AAS, JDO, SMS, JWR; Contributed to manuscript generation: AAS, JDO, SMS, JWR.

## Abstract

The principal mechanisms driving the synthesis and organization of durable animal structure have been the subject of intense investigation for decades. Here, we present evidence that mechanical strains can direct the formation of extracellular matrix (ECM) filaments via a mechanochemical cascade. This process is driven by cooperative cell contractions (“pulls”) that organize and precipitate structure via strain-induced polymer assembly. In an *in vitro* model of ECM synthesis, we use high-resolution optical microscopy to observe the kinematics of cell motion during their growth to confluency and identified cell-to-cell pulls that result in the production of persistent ECM filaments. Using live-cell confocal imaging, we confirmed that these pulls can directly cause the formation of fibronectin filaments that then bind collagen, producing persistent structures aligned with the direction of the pull. The finding suggests a new model for initial durable structure formation in animals based on local cell contraction, extensional strain and polymer mechanochemistry. The results have important implications for ECM development, growth and life-threatening pathologies of the ECM such as fibrosis.

## INTRODUCTION

The function of load-bearing (e.g. tendon) and other highly-specialized tissues (e.g. the cornea) is critically dependent on the precise placement and retention of structural polymers directly in the path of mechanical force. In vertebrates, fibroblastic cells work with their associated molecular secretome to efficiently generate a fully-continuous, mechanically-competent, polymeric extracellular matrix (ECM), orders of magnitude larger than themselves. The precise mechanisms driving ECM formation remain obscure because it is difficult to observe morphogenesis at high enough temporal and spatial resolution to determine the fate of structural molecules as they move from synthesis, to secretion, to incorporation into the developing durable tissue. Thus, most knowledge has been gained indirectly through destructive, static, high-spatial resolution electron microscopy ^1–6^ or lower spatial resolution optical microscopy ^5,6^. While dynamic live-imaging studies, precisely focused on ECM deposition, have the potential to reveal the mechanism of matrix formation, there have been very few of them ^7,8^ and they have not examined the correct temporal and spatial window to reveal the mechanism of deposition. We are thus left with little data capable of explaining structural causality with respect to the morphogenesis of organized connective tissue.

It is known that the formation of durable structure (e.g. collagen fibrils) depends on the formation of adhesion complexes on the cell surface ^9^, the formation of actin filaments ^10^, the presence of fibronectin (FN) ^11^, and the ability of cells to provide a force or to “pull” ^12–14^. In 2016, we demonstrated that extensional strain alone could drive collagen fibrillogenesis in a cell-free solution of collagen molecules ^15^. In this study, extensional strain rates on the order of 0.3 s^-1^ caused the flow-induced crystallization (FIC) of collagen molecules, a concept well-known in polymer physics ^16,17^. Flow-induced aggregation (FIA) of a number of important biomolecules (such as fibronectin and tau protein) has also been demonstrated ^18–22^. We suggested that fibroblasts might be capable of initiating assembly of ECM materials by pulling receptor-laden “filopodia” through an extracellular macromolecular milieu containing crowded ECM components, including collagen, fibronectin, and hyaluronan (HA) ^15^. Fibronectin is a particularly important player as it can “catalyze” collagen fibril formation during FIC ^20^.

Using an *in vitro* model of corneal stromal development ^8,23,24^, we investigated whether human fibroblasts use mechanical tension to polymerize ECM filaments. The model permits live, direct observation of primary human corneal fibroblast (PHCF) cell-matrix interactions. Using differential interference contrast (DIC) microscopy, we can take images at relatively high spatial (structures down to ~30 nm are detectable) and temporal (seconds between images) resolution. Here, we combine our PHCF ECM morphogenesis model with live DIC and confocal fluorescence imaging to examine ECM deposition during a critical window where durable matrix is initially formed.

### Live DIC Microscopy revealed that pericellular protrusions and filaments form via multiple types of cell contractions

PHCFs generate filaments and pericellular protrusions via rapid cellular contractions which we term “pulls”. The pulls were productive in that they “caused” the formation of persistent filaments. Five types of pulls were observed: 1) flat cell process (CP) pull; 2) thick CP pull; 3) thin CP pull; 4) ultrafine CP pull; and 5) cell surface pull. **Fig. 1** and **Supplemental Video 1** show representative pulls of each category along with plots of the measured velocities and extensional strain rates for that contraction. **Supplemental Fig. 1** and **Supplemental Video 2** show multiple examples of each pull type. In cases where two cells were involved, the pulls were further categorized as either symmetric (both cells contracted in what appeared to be a coordinated event) or asymmetric (only one cell contracted). When single-celled contractions occurred, the cell pulled against the culture dish surface and deposited filaments directly on the glass. **Supplemental Fig. 2** and **Supplemental Video 3** show representative still images and movies for the subcategories of symmetric, asymmetric, or single-celled pulling.

**Fig 1.**
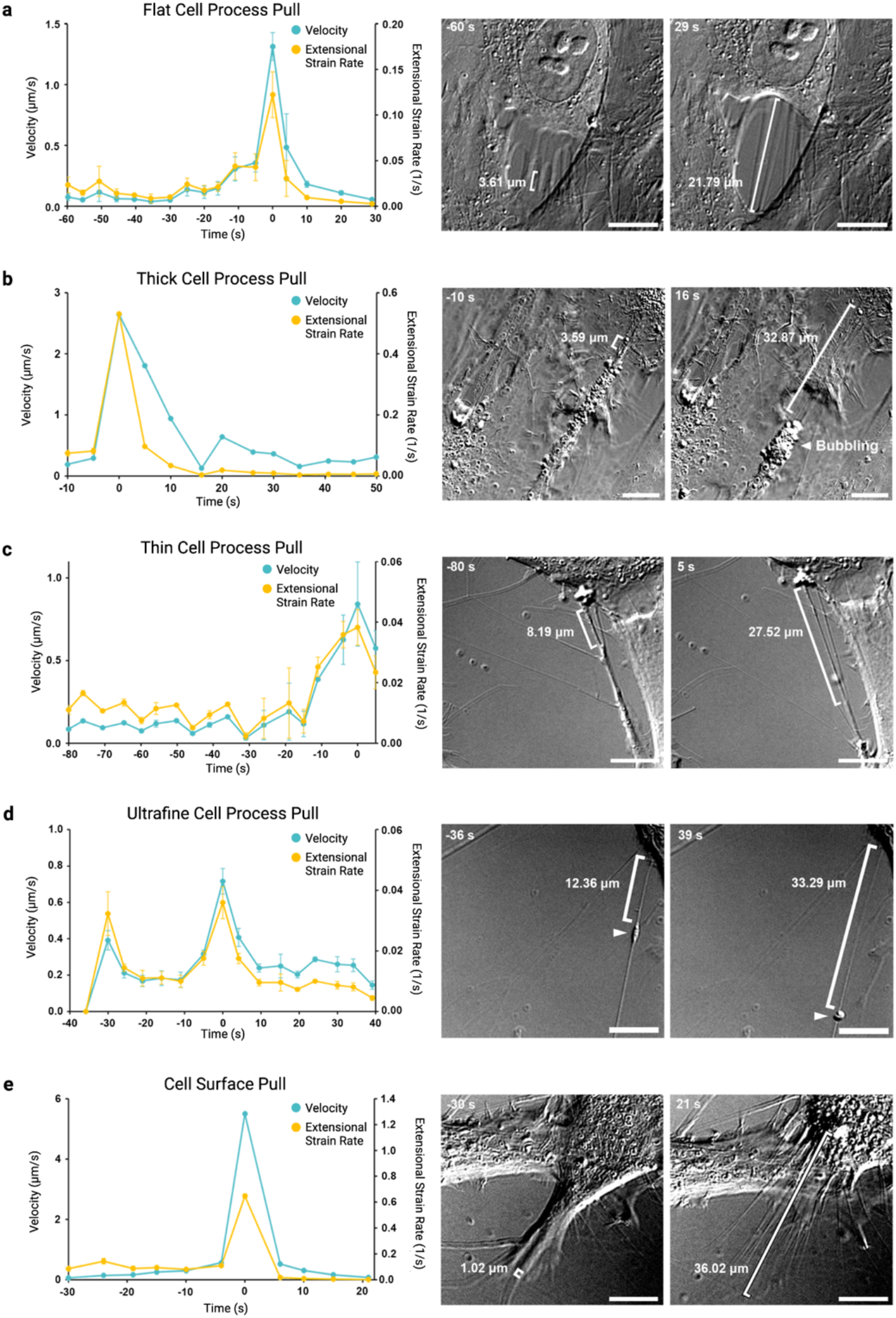
Five pull types with corresponding velocity and extensional strain rate plots. a-e, Representative PHCF DIC live-cell images of each pull type at initial and final timepoints (right), alongside their corresponding kinematics plots of velocity and extensional strain rate (left). Kinematics data is presented as mean ± standard deviation (n = 3 observers), while the filament (shown in brackets) is stretched during the pulling event. Generally, pulls exhibit a slow initial contraction, followed by a sharp rise to maximum velocity, and conclude with a sharp velocity decrease after the filament has been stretched. Flat CP pulls (a) are created by transparent lamellipodia-like projections on the edge of cells that create nearly parallel filaments aligned with the pulling direction and perpendicular to the cell border. Thick CP pulls (b) exhibit a wide “foot pad” at the end of the cell process which can pull to create many filaments. Nearly 90% of thick CPs appeared to be “bubbling,” or releasing vesicles into the extracellular space, at their foot pads. Thick CPs were also generally aligned with the cell’s long axis and often constitute the entire trailing edge of a cell. Thin CP pulls (c) are significantly thinner (p<0.05) than thick CP pulls, could be aligned at any angle relative to the cell long axis, and do not generally constitute the entire trailing edge of a cell. Nearly 50% exhibit minor bubbling, either by the CP itself or by the collaborating cell at the base of the forming filament. Ultrafine CP pulls (d) were created by the thinnest and longest CPs which were reminiscent of tunneling nanotubes, and most contained gondola-like vesicles (arrows) that were transported with the pull. Cell surface pulls (e) occur when the cell rapidly contracts its trailing edge as it migrates across the field of view, or when two adjacent, parallel cells rapidly separate. Cell surface filaments appear to originate directly from nucleation points on the cell surface as opposed to flat CP filaments which were formed by stretching the flat projection. Scale bars are 10 μm.

**Table 1** shows the results of our CP and filament analysis. 130 productive pulling events (i.e., produced a filament or filaments) were recorded and categorized based on pull characteristics. The number of filaments formed from each pull, filament maximum length, and total number of filaments analyzed in each category were recorded. We further assessed the length and thickness of each filament-producing CP.

**Table 1.** Summary of number of pulls, filaments formed, filament length, cell process thickness, and cell process length for each type.

| Pull Type | Number of Pulls Identified | Number of Filaments Formed Per Pull | Total Number of Filaments Analyzed | Average Filament Maximum Length (μm) | Cell Process Average Thickness (μm) | Cell Process Average Length (μm) |
| --- | --- | --- | --- | --- | --- | --- |
| Flat Cell Process | 50 | 4.8 ± 2.9 | 234 | 20.6 ± 11.3 | N/A | N/A |
| Thick Cell Process | 18 | 5.2 ± 3.0 | 94 | 21.8 ± 11.2 | 2.8 ± 0.7 | 41.9 ± 20.0 |
| Thin Cell Process | 21 | 3.8 ± 1.6 | 80 | 17.0 ± 10.0 | 1.3 ± 0.6 | 15.2 ± 7 |
| Ultrafine Cell Process | 5 | 1.2 ± 0.4 | 6 | 60.0 ± 21.7 | 1.0 ± 0.2 | 48.2 ± 14.8 |
| Cell Surface | 36 | 7.2 ± 4.7 | 253 | 26.4 ± 21.7 | N/A | N/A |

**Table 2** and **Fig. 2** summarize the kinematics data. Generally, pulls exhibited an initial slow contraction, then a sharp rise to maximum velocity, followed by a sharp decrease as the pulling event ended. The average maximum pull velocity was 0.80 ± 0.93 μm/s (n = 130) and the fastest pull was over 5 μm/s, or ~300 times the average speed of these cells on glass ^8^. This suggests that cell movement was not the primary objective of the pulls.

**Fig 2:**
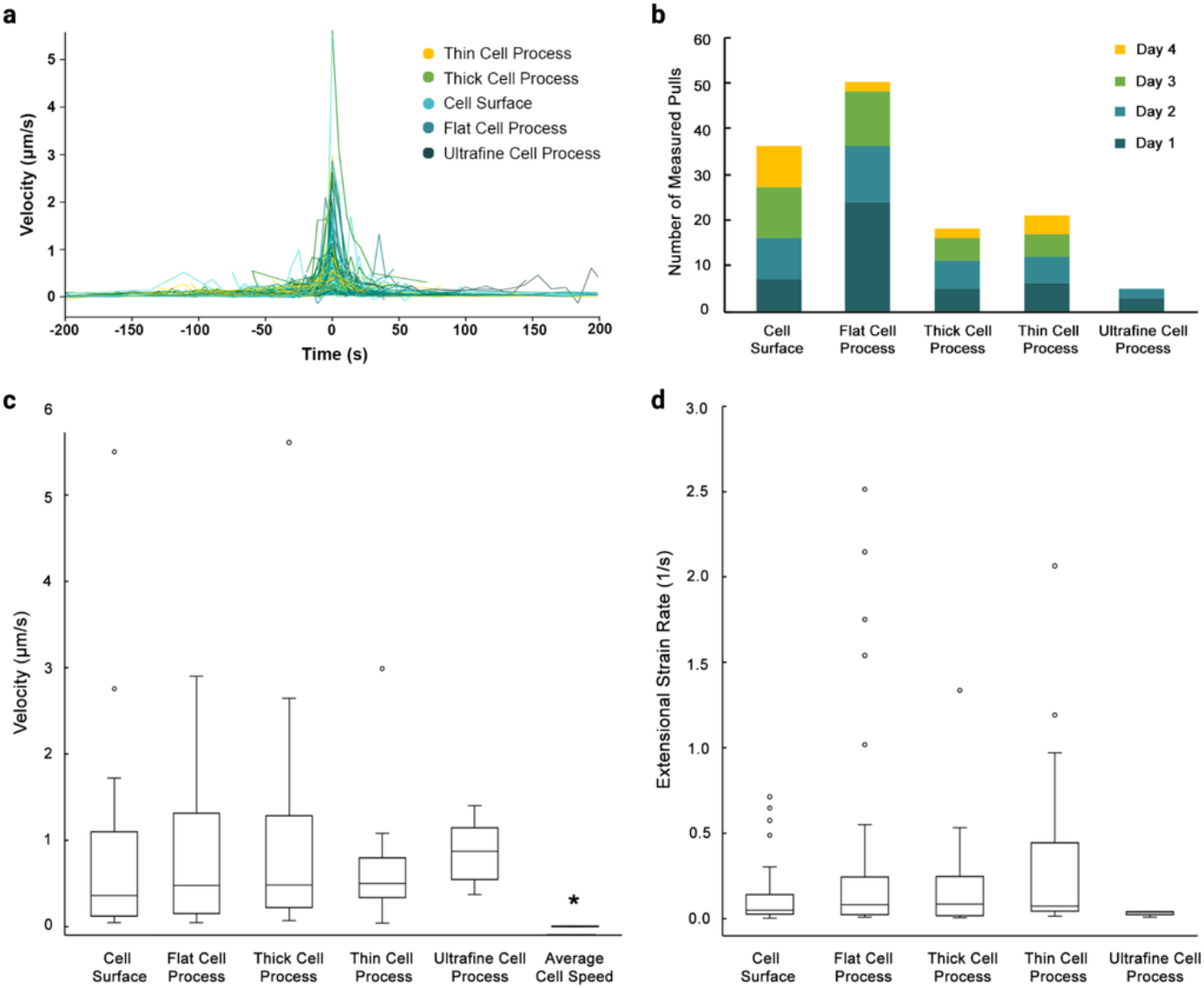
Pull kinematics data. a, Velocity profiles of assessed pulling events (n=130) standardized so maximum velocity is at time=0. Pulls typically begin slowly as the pulling elements begin to stretch their attachments and build tension in the structures that connect the pulling cell to a collaborating cell or to the glass (left side of the plot). Next, as more contractile elements engage and some structures yield to the applied load, the pull accelerates to a peak retraction velocity and extensional strain rate (center of plot). Finally, the pull sharply declines in speed (right side of plot) as the contractile elements bunch together, locally compacting the cell or cell process and halting the motion. b, Plot of the assessed pulling events, showing the distribution of events across days 1-4 and pull types. c, Box plot of the maximum pull velocity for each pull type. Average cell speed was significantly slower (p<0.01) than the maximum velocity of all pull types. d, Box plot of the maximum extensional strain rate for each pull type.

**Table 2.** Summary of average maximum pull velocities and extensional strain rates per pull type.

|  | Cell Surface | Flat Cell Process | Thick Cell Process | Thin Cell Process | Ultrafine Cell Process |
| --- | --- | --- | --- | --- | --- |
| **Total Contractions** | 36 | 50 | 18 | 21 | 5 |
| **Average Maximum Velocity ($\mu\text{m/s}$)** | $0.76 \pm 1.02$ | $0.82 \pm 0.82$ | $1.05 \pm 1.32$ | $0.64 \pm 0.59$ | $0.85 \pm 0.33$ |
| **Average Maximum Extensional Strain Rate ($\text{s}^{-1}$)** | $0.13 \pm 0.18$ | $0.30 \pm 0.55$ | $0.20 \pm 0.31$ | $0.33 \pm 0.51$ | $0.03 \pm 0.01$ |

The average maximum extensional strain rate was 0.23 ± 0.44 s^-1^ (n = 130) and the highest extensional strain rates created by flat CP, thick CP, and thin CP pulls exceeded 1 s^-1^. These strain rates are within the range (0.3 s^-1^) capable of forming collagen fibrils via FIC in a cell-free system ^15^. However, all pulls analyzed were “productive” in that they produced filaments, even at the lowest extensional strain rates of 0.01 s^-1^.

To determine if filaments were being pulled pre-formed out of a cell compartment or stretched, we assessed the thickness change of a subset of pulls using DIC microscopy ^25^ (n=9). **Supplemental Fig. 3** shows thinning of filaments during the contraction. However, we cannot state with any certainty that *all* filaments thinned during the pulling events; indeed, there appear to be cases where some filaments are formed from vesicular contents directly, and likely do not thin with deposition.

### Cell culture progression and pulling event observations

**Fig. 3** shows the progression of the cell culture by DIC and live-cell fluorescence microscopy over the four days of observation and at a later timepoint on day 7. While the cell density of the culture increases each day, it is clear from the live-labeling that filaments containing both collagen and fibronectin form at the earliest time points, with both biopolymers generally increasing in quantity over time.

**Fig. 3:**
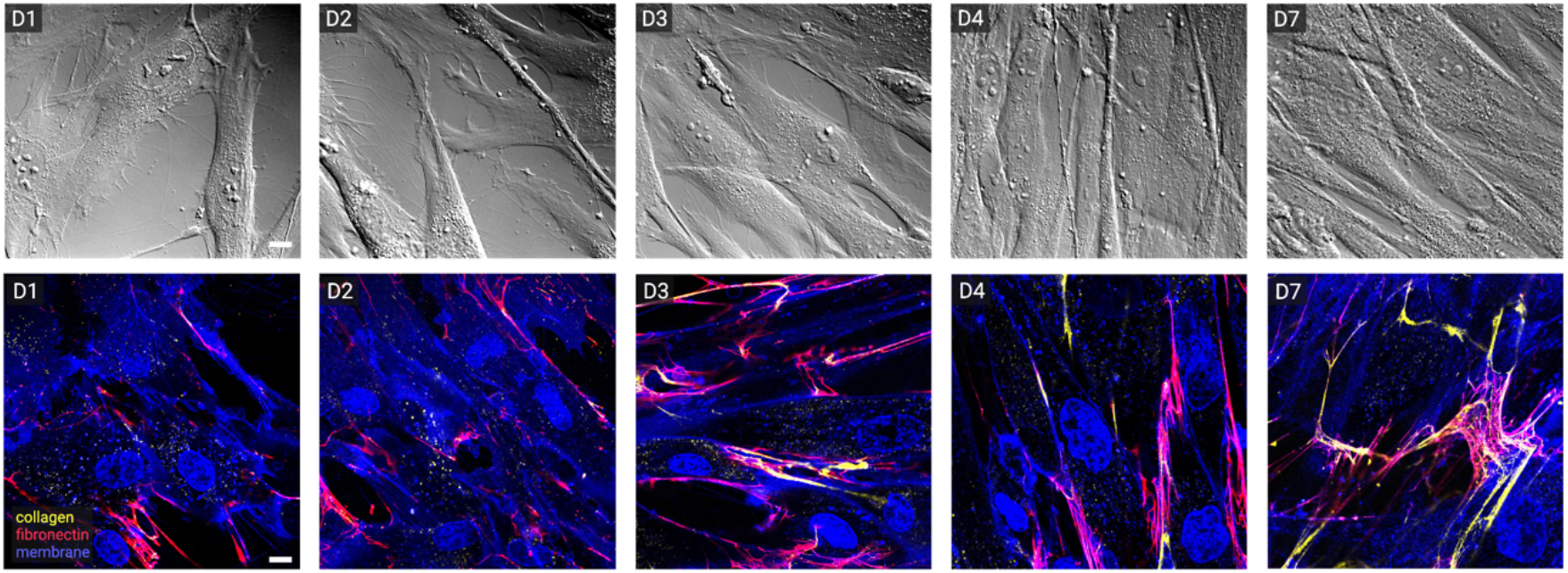
Cell culture progression. Representative DIC (top) and confocal (bottom) live-cell images of PHCF cultures on days 1-4 and day 7. Cells are stained for nucleus (blue), membrane (blue), fibronectin (red), and collagen (yellow). On days 1-2, the cells appear flat and relatively sparse. Fibronectin is visible on some portions of the cell membrane, and punctate collagen is incorporated into some fibronectin-membrane filaments (dashed circles). By days 3-4, the cells are nearly confluent. There is more fibronectin stretched between cells, and collagen appears more linear than punctate (arrows), suggesting collagen filaments are forming. By day 7, some collagen filaments have separated from the fibronectin-membrane filaments (triangular arrow) to form a durable structure that continues to be organized by pulling events. Scale bar is 10 μm, all images are the same size.

**Fig. 4** and **Supplemental Video 4** show numerous events that may be of importance in interpreting the mechanisms in operation during filament formation. <u>Surface vesicles</u>. Previous live imaging data shows that cells exhibit surface “bubbles” (vesicles) in locations where filaments form ^26,27^. Bubbles were often observed at the site of filament formation immediately prior to the pull, and in some cases, they would shrink or disappear following the contraction suggesting that their contents may have been ejected or unspooled during the pulling event. We were also able to closely observe the direct deposition of filaments on the glass surface. <u>Bidirectional pulls</u>. There were numerous cases observed where the same cell would pull in one direction (typically along its principal axis) and then immediately pull at a different angle to the previous pull. In some cases, the angle was close to orthogonal, suggesting that there might be some internal bidirectionality or bi-polarization instrumental to the ultimate organization of the tissue being produced. In this culture system ^23^ and in their natural state, PHCFs produce nearly-orthogonal sheets of collagen. <u>Filament Reel-in</u>. In many pulling events, the filaments were not only extended by the retraction of the cell process that created them, but they were also “reeled in” at the same time. This likely makes the actual extensional strain rate of the filament higher than could be calculated by simply measuring the change in length of the filament.

**Fig 4.**
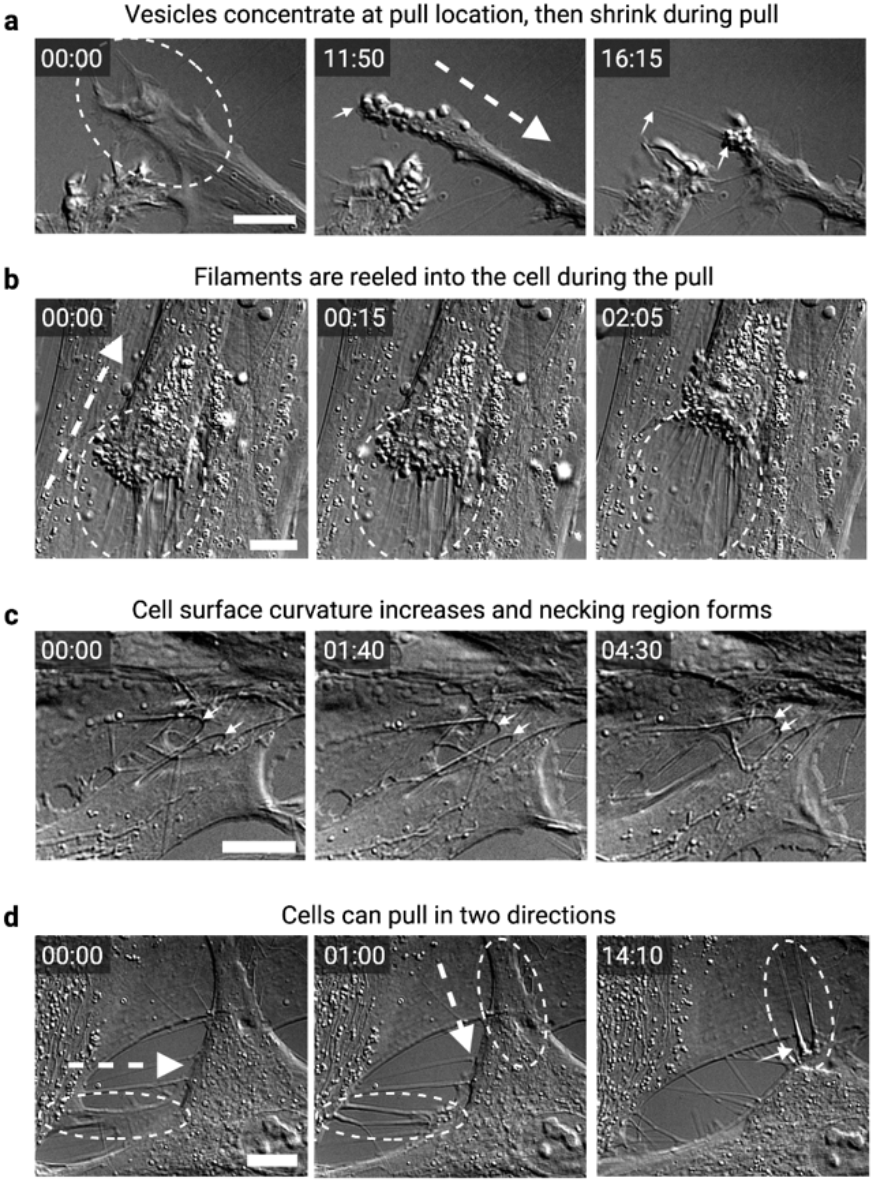
Notable observations that occur during pulling events. a-d, Live-cell DIC images of PHCFs showing examples of notable observations that occur during pulling events. a, Long before the pulling event (left), no bubbles are present. Immediately before the pulling event (middle) many spherical bubbles, or vesicles, appear and concentrate at the edges of the CP (indicated by the small arrow). As the cell pulled, a filament was deposited directly onto the glass (indicated by small arrows) and the bubbles shrunk and disappeared. b, During this pulling event, many bubbles form and concentrate at the cell surface where filaments are forming. As vesicles are depleted, new ones appear in a vigorous bubbling manner, and the cell appears to reel in the newly-created filaments. c, Before the pull, the curvature at the base of the filament (indicated with small arrows) is more open. As the pull occurs, the cell surface appears to become more curved, similar to a necking region. d, First, the cell’s left CP pulls inward and forms filaments. Next, the cell’s top CP pulls inward to form additional, nearly perpendicular filaments. These filaments were further reeled in, causing a crumpled region at the base of the filament (small arrow). Scale bars 10 μm, large dashed arrow indicates pull direction, and time is min:sec.

**Fig. 5 and Supplemental Video 5** show a clear example of flow aggregation and fibronectin/collagen co-association during a pulling event. In the video, we see the exportation of collagen to the cell surface followed by a pulling event that creates an array of continuous fibronectin filaments which bind to the exported collagen, pulling it along with the fibronectin. The fibronectin/collagen association is consistent across our observation period with nearly all observed collagen filaments colocalized to extending fibronectin filaments.

**Figure 5:**
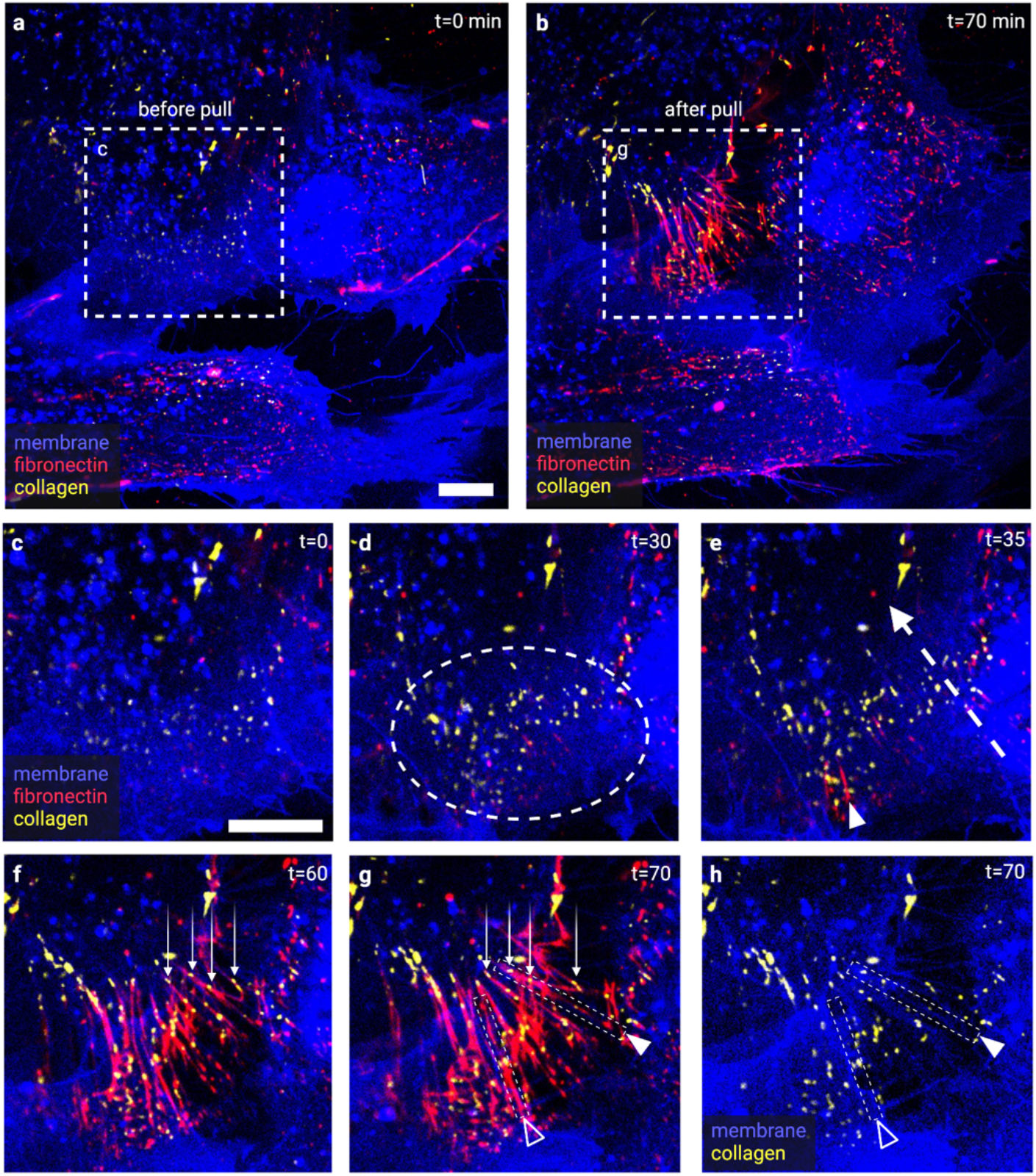
Pull causes ECM structure formation. Live-cell confocal images of a Day 1 PHCF culture stained for nucleus (blue), membrane (blue), fibronectin (red), and collagen (yellow). Images were taken of the underside of the cell, focused between the cell and the glass. a-b, Low magnification images before and after the slow cell surface pulling event causes ECM formation. c-e, High magnification inset showcasing the pulling event. Before the pulling event (c-d), punctate collagen and fibronectin globs (circled) become brighter and more abundant. As the pull begins (e), fibronectin appears to “streak” (small arrow) in the direction of the pull (large arrow). Fibronectin becomes brighter and begins forming a matrix of filaments (f). Collagen colocalizes with fibronectin and elongates with the stretching ECM (g). Arrows point to globs of collagen that move with fibronectin. h, By subtracting the fibronectin and enhancing the membrane, we can see a network of membranous cell filaments colocalized with most of the newly-formed ECM. g-h, White triangle points to ECM filament that has a membrane filament at its core (small rectangle), and hollow triangle points to an ECM filament that does not have a membrane core (small rectangle). Scale bars are 10 μm.

To determine if the same kind of pulling kinematics occurs in a different species, we examined the behavior of rat fibroblasts under the same conditions and saw similar filament formation and kinematic behavior (**Supplemental Fig. 4**).

### The effect of molecular crowding with Ficoll

Molecular crowding of corneal fibroblasts can convert secreted pro-collagen to telocollagen (i.e., activated for assembly ^28^) and drastically enhance the rate of matrix deposition ^29,30^. We conducted this experiment with and without Ficoll as a molecular crowding agent. While the addition of Ficoll to the culture system resulted in thicker constructs at two weeks 20.38 ± 5.81 μm vs 12.95 ± 1.71 μm (p<0.05), we did not observe significant differences in the types of pulls, maximum pull velocities, or maximum extensional strain rates during the initial four-day observation period in the crowded versus uncrowded experiments and thus pooled the data.

### Availability of the raw data

The data set represents an extensive resource that is likely to be of interest to other researchers in the field of ECM development and cell behavior. We will thus make the raw videos, in NIS format, available to anyone who would like to examine them upon request. Please place an acknowledgment that the videos were provided courtesy of Alexandra A. Silverman.

## DISCUSSION

Creating highly-anisotropic arrays of collagen fibrils is critical to the mechanical function of tensile load-bearing (e.g. tendon) and specialized connective (e.g. the cornea) tissues. Fibroblast cells possess contractile machinery and polarization and thus have a ready mechanism to drive and control ECM organization: mechanical force. Mechanical forces are vectors, with direction, and magnitude and which operate over long distances capable of controlling connective tissue deposition ^31^. Here we have demonstrated that persistent filamentous structures form secondary to the act of a fibroblast, or portion thereof, pulling towards itself. These pulls could involve single or multiple cells and different structures associated with each cell. Some of the pulls captured were of the highest velocities recorded in the literature for cell process retraction ^32^ and some caused extensional strain rates well above those shown to produce collagen fibrils *in vitro* from cell-free, pure collagen solutions ^15^. The fact that filaments also formed at very low strain rates has substantial implications for the underlying biophysics of durable matrix assembly and more detailed knowledge of the molecular composition of the filaments, the proximal source of the molecules involved, and the distribution of strain on the forming filaments is required. Our observations, performed at high spatial and temporal resolution support the principal hypothesis that structures (filaments) are formed preferentially “along the lines of local mechanical strain.” More succinctly, we have directly observed mechanical causation of biological structure at the cellular length scale. The active elements involved in the process appear to include contracting cell surfaces, cell surface protrusions, lamellipodia, cell processes and mechanosensitive biopolymers: fibronectin and collagen, at least.

### Pulls are discrete events with characteristic kinematics capable of flow-induced protein aggregation or crystallization of biological polymers

We label the events described in this study “pulls” to distinguish them from the normal movements of the cells. A pull is a rapid kinematic event that has a clearly characteristic motion comprising three parts: 1) Slow initiation, stretching attachments and building tension in the structures that connect the pulling cell to a collaborating cell or to the glass; 2) Acceleration to a peak retraction velocity, well above average cell velocity, and attainment of peak extensional strain rate; 3) Sharp deceleration. Only one subset of pulls (ultrafine), which we believe were generating tunneling nanotubes, did not follow this pattern. For four of the pull categories, flat CP, thick CP, thin CP, and cell surface pulls, the maximum extensional strain rates were sufficient to cause collagen fibril formation via FIC based on the data of Paten *et al*. (2016) ^15^ and well above that needed for hybrid collagen/fibronectin fiber formation via combined FIA/FIC as shown in Paten *et al*. (2019) ^20^.

### ALL filaments observed during their production were derivative of pulling events between filament “anchor” points

We did *not* observe any long filaments spontaneously forming in the culture system without an associated contraction event. Nor did we observe the pure extrusion of long filaments from cell invaginations or long filaments emerging from the cell surface in the absence of extensional mechanical assistance. All forming filaments started and ended at fixed points, either on a cell surface, a CP or on the glass. The initial connection between the surfaces that subsequently “separate” is thus an important mechanism to examine. Our data further suggest that filaments are not being pre-formed within the cell and drawn out. In many cases, filaments grew to lengths longer than the cell body, providing further evidence that the filament was assembling, growing, or stretching as a direct result of the pull. Cooperation between cells is likely critical to filament formation. Data on collagen fibril synthesis support this idea. Lu *et al*. (2018) reported that individual collagen fibrils are formed by more than one cell when they used co-cultures of two different colored collagen expressing cells ^7^.

We also observed that some filaments are drawn back into the cell process or “reeled in’’. While our data clearly demonstrate that filaments are formed extracellularly, we suspect that 1) some filaments are lengthened by being reeled in by cell processes and 2) in some cases if filaments break from one of their anchor points, they will be phagocytosed after being reeled in.

### Filament type and composition

Because of their fragility during fixation and immunohistochemistry staining (**Supplemental Fig. 5**), live cell stains were necessary to identify components within the filaments. We found that a majority of pulling events formed thin, membrane extensions, which were often the filaments observed during DIC imaging. We further found that collagen and fibronectin frequently bound to and stretched with these membrane extensions. Even at the earliest culture time points, collagen and fibronectin were present in pulled filaments, and collagen was clearly bound to and translocated with the fibronectin. These observations suggest that exported “globs” of collagen are captured and reformed into linear structures by association with the fibronectin.

Our findings are consistent with Young *et al*. ^4^ and Bueno *et al*. ^33^ who both observed that thin cell extensions associate with the forming collagen matrix. Young *et al*. termed these matrix-forming structures “keratopodia”. These cell extensions are likely important drivers of early matrix assembly and are possibly critical to fibrotic tissue formation where the cells are muscular and highly contractile.

### Proposed models of ECM filament formation

Based on our observations, we propose two possible models, along with live-cell imaging examples, of ECM filament formation driven by cellular contractions (**Fig. 6**). The molecules included in our models all are known to assemble into fibers, with fibronectin and collagen known to assemble under extensional strain ^15,19,34^. Additionally, it is known that pro-collagen is exported to the surface in secretory granules ^35^ and that it (and its pro-peptidases) is potentially bound to fibronectin to facilitate local activation ^36^. While we did not stain for HA, we include in the model as it is thought to assist in fibronectin/collagen deposition ^37^ and it is known that HA is synthesized by HAS-2 ^38^ near the membrane. Because all filaments were formed by the mechanical separation of two structures, both of our models begin with the formation of an attachment between cells.

**Figure 6:**
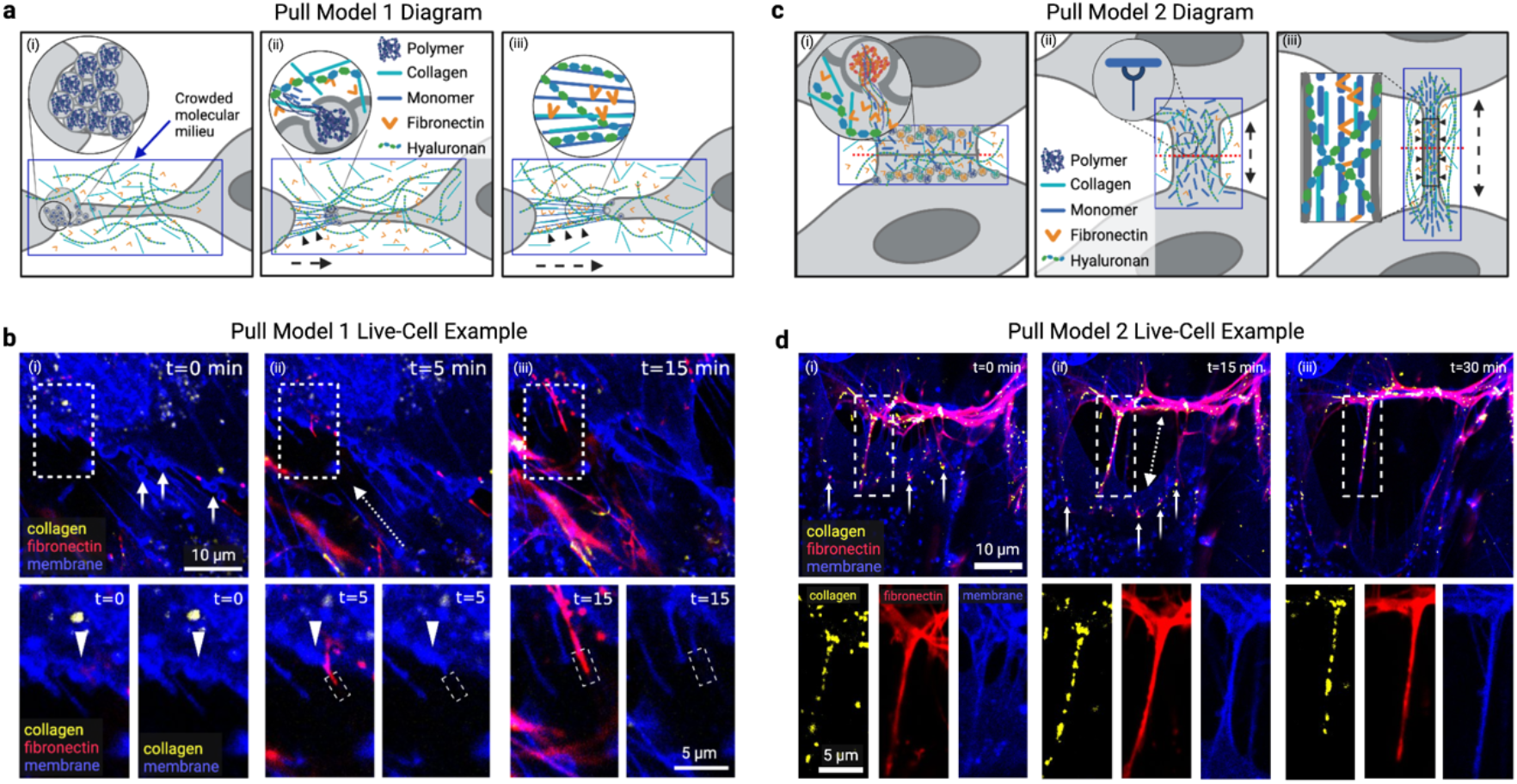
Two proposed models for mechanically driven filament assembly. Two proposed models (a, c) and corresponding live-cell confocal microscopy examples (b, d) for mechanically driven filament assembly. Confocal time-lapse images are of PHCFs stained for nucleus (blue), membrane (blue), fibronectin (red), and collagen (yellow). a, Model 1 shows filament formation without a cell membrane extension at the core. A bubbling cell process pulls (i) and releases biopolymer precursor molecules that become stretched and rapidly form filaments (ii). Then, additional biopolymers, or ECM components (collagen, fibronectin, hyaluronan), can participate in filament formation (iii). b, Live-cell example of Model 1. i, Cell membrane is bubbling (small arrows) and minimal fibronectin is present. ii, As the cell pulls (direction indicated by dashed arrow), a fibronectin filament streaks out of a vesicle (triangle). The end section of the filament (dashed rectangle in inset) comprises only fibronectin and no membrane. iii, Filament grows and the section comprising only fibronectin becomes longer and thicker. c, Model 2 shows filament formation with a cell membrane extension at the core. i, Two adjacent cells export biopolymer precursor molecules to the surface and release them at the site of the pull. ii, The molecules bind to the cell surface and the cells begin to separate, forming a membrane bridge. iii, As the cells pull, the membrane bridge thins and elongates, causing the bound molecules to align, concentrate, and extend along the path of load, leading to ECM polymer crystallization. d, Live-cell example of Model 2. i, A filament comprising membrane, fibronectin, and collagen is stretched between two adjacent cells. Globs of fibronectin and collagen (small arrows) are present on the cell surface. ii-iii, As the cell pulls (direction indicated by dashed arrow), the filament becomes longer and thinner.

#### Model 1: Pure FIA/FIC

**Fig. 6 a-b** shows filaments that are generated without a cell membrane extension at their core. The attachment point is formed such that biopolymer precursor molecules are positioned between the two surfaces. **Fig. 6 a-b, i** shows precursor packed into vesicles at the attachment point. Upon pulling, the dense polymer solution is stretched, rapidly forming filaments. **Fig. 6 a-b, ii** shows the initial stretch and the polymers “spooling” out of the vesicles as filaments. As the stretching progresses (**Fig. 6 a-b, iii**), the filament lengthens and the dense molecular milieu outside the cells (pre-synthesized or concurrently ejected from vesicles) may participate in the filament composition. The first pull in **Supplemental Video 4** where the cell deposits matrix directly on glass is a clear example of model 1.

#### Model 2: Cell Process/Receptor Mediated FIA/FIC

**Fig. 6 c-d** depicts a stretching filament which has a long thin cell surface extension at its center. In this model, the cellular contraction stretches the cell membrane(s) into a long, thin filament, replete with cell surface receptors that bind extracellular structural biopolymers (pre-synthesized or concurrently ejected from vesicles, **Fig. 6 c-d, i-ii**). This reduces biopolymer motility and increases their effective relaxation time. As the pull continues, the filament thins and lengthens, aligning the bound polymers, extending them in the direction of load, and concentrating them circumferentially (**Fig 6 c-d, iii**). This produces physical conditions which, we hypothesize, promote the crystallization of the bound polymers.

All three of our candidate molecules are ligands for cell receptors found on corneal and other fibroblasts (HA-CD44 ^39,40^; FN:a5b1 ^41^; and Col:a2b1 ^42^). Additionally, it has been demonstrated that monomeric collagen in solution rapidly homes to and incorporates with collagen fibrils under strain ^43^. Model 2 is also consistent with the co-alignment of corneal cell processes (“keratopodia”) with collagen arrays observed by Young *et al*. ^4^ and Bueno *et al*. ^33^.

In both Model 1 and Model 2, the durable filaments (collagen) separate from the fibronectin after it facilitates their formation, completing the process of initial matrix morphogenesis (**Supplemental Fig. 6**).

### Limitations of the experiment

First, there is limited physiological relevance of early cultures on glass. Our experimental model is a human primary cell culture system that is known to produce orthogonal arrays of collagen, in an arrangement similar to the native cornea stroma, over a period of weeks. Early on, the cells cannot produce extensive matrix because collagen is lost to the media prior to conversion by propeptidases ^44^. Additionally, these cells behave more like myofibroblasts than fibroblasts in early culture. However, prior to confluence, we can observe and quantify their behavior with high-resolution DIC. Lastly, there are potentially interfering effects of fibronectin and other components in the system due to the presence of bovine serum in the media.

### Implications of the observed behavior

There are a number of potential implications for ECM development, growth, maintenance, repair, and pathology if we cautiously extend the fundamental pulling behaviors observed in our model system to the *in vivo* cell-ECM relationship.

#### A new model of durable ECM formation: Mechanochemical Force-Structure Causality

The first implication is the potential discovery of a fundamental, proximal mechanism driving organized durable ECM production, which we posit to be mechanochemical in nature. In all cases, we observed that pulling produced the filaments. We describe this activity as *mechanical causation of structure*, or more formally, mechanochemical force-structure causality (MFSC). Mechanical strain has previously been shown to *cause* structure formation for two of the three major structural biopolymers involved in the development of ECM: fibronectin ^45^ and collagen ^15^. In the case of fibronectin, the deformation of the molecules opens a cryptic binding site to promote fibronectin fiber assembly via FIA ^45^. For collagen, extensional strain-rates align molecules accelerating assembly via FIC ^15^. When fibronectin and collagen are together, there is a synergistic effect lowering the energy barrier to assembly further ^20^. The most important aspect of MFSC is that structure is formed directly in the path of the forces that create it, ostensibly to resist subsequent forces which propagate along the same direction. This is possibly why we first see cell and cell process alignment with the eventual matrix that is formed in developing organized tissues ^1,4,46,47^. Cells produce the initial structure along local lines of “pull” and then global level tissue forces (i.e., muscle contractions, growth pressure increases) complete the structure formation, as we described in Paten *et al*. (2016) ^15^. It has been shown that knocking out non-muscle myosin II disrupts initial collagen fibril formation ^13^, abrogating muscle contractions cause tendon formation to fail ^48–51^, and de-pressuring the ocular globe leads to impaired growth ^52^, underscoring the importance of cell-generated forces.

#### Prestress: Filament Deposition under Tension

Tissues generally exhibit prestress, suggesting fibril deposition under tension ^53^. If mechanical strains are required to cause the formation of structure, then the structures that are formed are going to be pre-strained by design. It is interesting to note that the seminal demonstration of engineered tissue formation was a collagen gel contracted by fibroblasts ^54^.

#### Substrate Effective Stiffness: Soft vs. Hard Substrates

Our cells were plated on glass which is, in effect, infinitely stiff. Thus, cells can generate maximum forces and velocities, provided they were well-anchored to the substrate. On very soft materials, fibroblasts cannot generate the same separation forces or speeds, limiting their ability to produce structure. Fibroblasts on soft tissues under load would experience an effectively higher resistance in the direction of force and strain, and therefore would likely be more effective at producing structure precisely where the loads are highest.

#### The Structural Organization Kernel: Effect of Cell Polarization

In our system, we used corneal stromal cells which inherently produce orthogonal layers of parallel collagen fibrils. We found many examples of double pulls where cells would pull in one direction and then follow that quickly by pulling in another direction, which was often close to orthogonal. Electron micrographs ^1^ and reconstructions ^4^ of developing chick cornea show that the same cell has collagen fibrils extending in two directions from the cell body. Therefore, multiple pulling directions could be fundamental to the ultimate organization of the tissue being constructed.

#### Fibrosis, Wound Healing, and Tissue Engineering

Nearly 45% of all deaths worldwide are related to some form of fibrosis ^55^. It is well-known that increased contractile machinery is present in myofibroblasts and that the transformation of local fibroblasts to myofibroblasts is a marker of fibrosis ^56^. The increased muscularity of myofibroblasts would cause the formation of pathogenic structure through MFSC, provided there were available precursor biopolymers. Further, as structure is produced and stiffens, the effectiveness of each pull increases, leading to more accumulation of ECM in a positive feedback loop. Once we understand the incipient mechanisms of fibrosis, there are opportunities to control or reverse it. As a corollary, wound healing and engineering of ECM tissues could be enhanced through the harnessing of MFSC principles.

## Conclusion

Our data suggest that local pulling leads to durable structure formation in an *in vitro* model of developing ECM. The nature of the pulling profile indicates that pulling events are separate and distinct from those that provide cell motility or apply tension in general. Tying mechanical force and the resulting strain rates directly to biopolymer structure formation via FIA and FIC could lead to a clearer, functional understanding of the cell-matrix interplay during development, growth, and pathological changes of connective tissues (e.g., fibrosis). There are rather important implications associated with the proposed MFSC model. However, on a more philosophical note, if millions of micron-scale, cooperative cell contractions cause the formation of tiny polymeric strings that integrate to form our load-bearing structure, then we could be, quite literally, “pulled” into existence.

## MATERIALS AND METHODS

### Cell Isolation and Culture

PHCFs were isolated from 74-year-old human donor corneas (obtained through the National Disease Research Interchange) following the procedure established by Bueno *et al*. (2009) ^33^. Briefly, donor corneas were gently scraped of their epithelium and endothelium to ensure only stromal cells were isolated. Then, the corneas were cut into 2×2 mm pieces and sterilized by soaking in 1X Gibco phosphate buffered saline (PBS) (Fisher Scientific, 10-010-023) containing 1% HyClone penicillin streptomycin 100X solution (Fisher Scientific, SV30010) and 0.1% amphotericin B 250 μg/mL solution (Sigma Aldrich, 1397-89-3). Each explant was adhered to the center of a 6-well culture plate (Corning, 3516) and PHCF were cultured using the reagents and techniques established by Siadat *et al*. (2021) ^57^. Complete media, comprising Corning DMEM with L-glutamine and 4.5 g/L glucose without sodium pyruvate (Fisher Scientific, MT10017CV), 1% penicillin streptomycin, 0.1% amphotericin B, and 10% Corning regular fetal bovine serum (Fisher Scientific, MT35010CV), was gently added to the well. Cells were incubated at 37 *°C* with 5% CO_2_. A half media exchange was performed every 3-5 days until the PHCF migrated off the explant. Once confluent, the explants were discarded and PHCF were expanded and frozen for future use.

### Live-Cell DIC Imaging

PHCF were seeded on uncoated, 1.5 thickness glass-bottomed, temperature-controlled culture dishes (Bioptechs, 04200417B) at passage 3 at a concentration of 10,000 cells/cm^2^. On day 1, a full media exchange was performed and CO_2_-infused, complete media containing 0.5 mM L-ascorbic acid (Sigma Aldrich, A4544) was added to stabilize the collagen produced by the culture. For half the samples, 37.5 mg/mL Ficoll 70 (Sigma Aldrich, F2878) and 25 mg/mL Ficoll 400 (Sigma Aldrich, F8016) was added to the media to increase the molecular crowding of ECM proteins. 6 samples treated with only ascorbic acid and 6 samples treated with ascorbic acid, Ficoll 70, and Ficoll 400 were assessed.

After the media was exchanged, the cells were incubated for 3 hours and then imaged with a coverglass lid (Bioptechs, 4200312) using a Nikon ECLIPSE TE2000-E inverted microscope. A 60X Nikon CFI Apochromat TIRF oil objective and Photometrics CoolSNAP EZ CCD camera was used to take high-resolution, DIC photos every 5 seconds for one hour. The Nikon perfect focus system was utilized to ensure the sample remained in focus during imaging. To keep the sample at 37 °C, an objective heater (Bioptechs, 150819), a microscope stage adapter (Bioptechs, 04202602), an objective heater controller (Bioptechs, 150803), and a culture dish temperature controller (Bioptechs, 0420-04-03) were used. All samples were imaged at approximately the same location to ensure a consistent distribution of cells.

After imaging the sample, it was placed back in the incubator to be subsequently imaged on day 2, 3, and 4. On days 2 and 4, no media exchange was performed prior to imaging. On day 3, a full media exchange was performed 3 hours prior to imaging using CO2-infused complete media containing either only 0.5 mM L-ascorbic acid or 0.5 mM L-ascorbic acid, 37.5 mg/mL Ficoll 70, and 25 mg/mL Ficoll 400.

### Live-Cell Confocal Imaging

CNA35-GFP, a collagen binding protein with green fluorescent protein, was prepared following the protocol established by Aper *et al^58^*. Briefly, single *E. coli* colonies with the pET28a-EGFP-CNA35 gene (Addgene, Plasmid #61603) were cultured in LB media and protein expression was induced with 0.5 mM isopropyl β-D-1-thiogalactopyranoside (Sigma Aldrich, I1284). Bacteria pellets were resuspended in Bugbuster (Sigma Aldrich, 70584-3) containing 1 μL/mL benzonase (Fisher Sceintific, 70-746-3). Protein was purified via Ni2+-affinity chromatography using the N-terminal 6xHis-tag.

Fibronectin stain was prepared by mixing 100 μL Alexa Fluor 594 anti-fibronectin antibody (Abcam, ab275336) with 0.3 mL glycerol (Fisher Scientific, G33), 0.01 g bovine serum albumin (Sigma Aldrich, A8654), and 0.6 mL 1X phosphate buffered saline (Fisher Scientific, AAJ67670AP) to create a stock solution of 50 μg/mL antibody which was stored at −20°C until use.

For each sample, both nucleus-membrane and collagen-fibronectin staining solutions were prepared with complete media containing 0.5 mM L-ascorbic acid, 37.5 mg/mL Ficoll 70, and 25 mg/mL Ficoll 400. Nucleus-membrane staining media was supplemented with 2 drops/mL of NucBlue (ThermoFisher, R37605) and 1 μL/mL CellBrite Steady Membrane Dye (Biotium, 30105-T and 30108-T). Initially, CellBrite Steady Membrane Dye 405 nm was used, but it was replaced with CellBrite Steady Membrane Dye 650 nm due to phototoxicity. Collagen-fibronectin staining media was supplemented with 20 μL/mL CNA35-EGFP and 15 μL/mL fibronectin antibody.

Cells were cultured on Bioptechs culture dishes with complete media containing 0.5 mM L-ascorbic acid, 37.5 mg/mL Ficoll 70, and 25 mg/mL Ficoll 400 as previously described. Prior to imaging, a full media exchange was performed and 1 mL of nucleus-membrane staining media was added and cells were and incubated for 30 minutes at 37°C and 5% CO2. Then, another full media exchange was performed and 1 mL of collagen-fibronectin staining media was added and cells were incubated for at least 30 minutes at 37°C and 5% CO2. This collagen-fibronectin staining media was kept in the dish during all confocal imaging.

Cells were imaged on a Zeiss LSM 880 confocal laser scanning microscope using a 63x oil objective with controlled incubation settings of 37°C, 5% CO2 and XX% humidity. Cells were imaged using 405 nm, 488nm, 561 nm, and 640 nm laser lines with a pixel dwell time of either 0.55 μs or 1.1 μs. The laser intensity, pinhole size, and gain were optimized to decrease photobleaching while enabling visualization of fine filaments. Live-cell confocal time lapse images were taken every 5 minutes. Zstacks and tile scans were also performed. After imaging, cells were discarded.

### Assessing Cell Culture Thickness

After imaging on day 4, samples were grown until day 14 with full media exchanges every 2-3 days using complete media with consistent concentrations of either only L-ascorbic acid or L-ascorbic acid, Ficoll 70, and Ficoll 400. On day 14, cell construct thickness was assessed by using the Nikon inverted microscope with the 60X oil objective and DIC to perform a Z-scan in center, north, south, east, and west locations in each cell culture. Multiple locations were used due to heterogeneity in the cell culture; these locations were averaged to obtain an average thickness for each cell construct.

### DIC Time-Lapse Analysis

Time-lapse videos were analyzed frame-by-frame using FIJI. For each identified pull, the length of the persistent filament formed was measured over time, and the following equations were used to analyze the pulling events:

PHCF pull velocity: 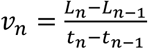

PHCF pull extensional strain rate: 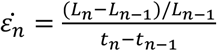

Where *L_n_* is the length of the persistent filament in frame *n*, *L_n-1_* is the length of the persistent filament in the previous frame (*n*-1), *t_n_* is the time in frame *n*, and *t_n-1_* is the time at the previous frame. Though our microscope was set to take photos every 5 seconds, this timing was not always precise, and images were taken approximately every 4-6 seconds. For our analysis, we used the exact time points exported from our data.

A total of 130 pulling events were analyzed, individually by three authors to obtain 3 measurements for length, 3 measurements for velocity, and 3 measurements for extensional strain rate for each pull at each time point. These values were averaged to obtain a mean length, velocity, and extensional strain rate for each pull at each time point. The maximum velocity and extensional strain rate for each pulling event was used to compare events.

Pulling events were subsequently categorized into five pull types based on pull morphology. For pulling events that created multiple persistent filaments, we recorded the total number of filaments created and their maximum length, and then analyzed the most prominent and visible filament. The length of the contracting cell process and its average thickness was recorded. To determine cell process thickness, measurements were taken at 5 locations along the process and then averaged. For each pulling event, it was noted if the cell process contained a bubbling footpad that released vesicles into the extracellular space. It was also recorded whether the event occurred between two cells, or one cell and the glass surface of the culture dish.

### DIC-EIS Analysis

Filament diameter change was estimated from DIC images based on a method that measures collagen fibril diameter ^25^. Only filaments that stayed in focus during the entire pulling event were analyzed. DIC images were uploaded into Matlab. A rectangular region of interest was defined along the middle section of each filament. DIC-EIS was calculated as the average difference between the maximum and minimum intensities across each filament.

### Video Creation

Files were imported into FIJI as an image sequence and cropped, rotated, and adjusted to have increased contrast and better showcase the pulling event. The scale bar and time stamp were also added in FIJI. Our time stamp was set to increase in increments of either 5 seconds (for DIC imaging) or 5 minutes (for confocal imaging). Photoshop and Biorender were utilized to create video annotations and iMovie was used to assemble and edit the videos.

## Acknowledgments

This work was supported by funding from NIH/NEI 1R21EY029167. Images created with Biorender.com. We thank the Institute for Chemical Imaging of Living Systems at Northeastern University for consultation and imaging support.

## Supplemental Figures

**Supplemental Fig 1.**
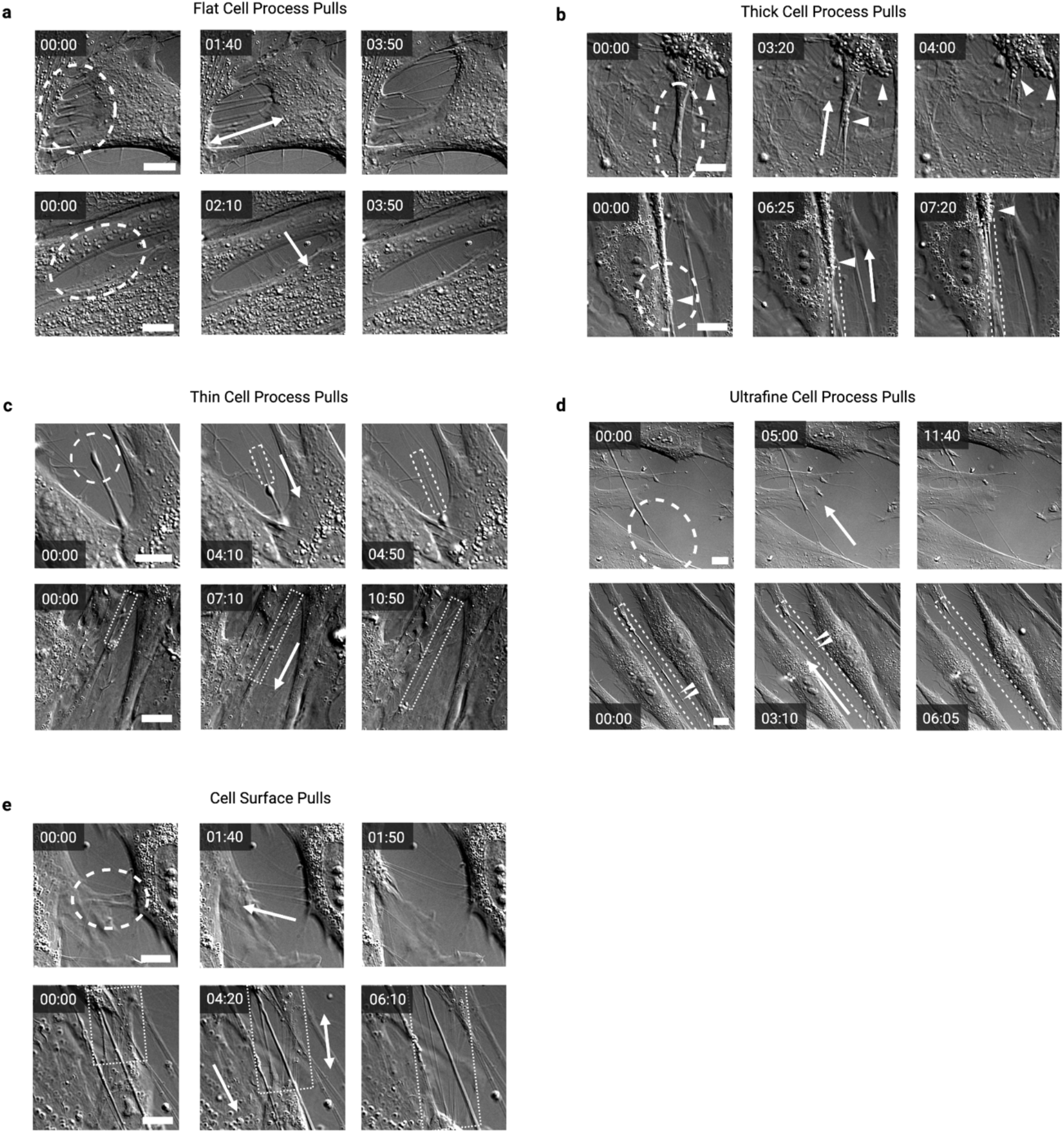
Additional examples of each pull type. a-e, PHCF DIC live-cell images of two additional examples for each pull type. Flat CP pulls (a, n=50) are created by transparent lamellipodia-like projections on the edges of cells. These pulls can occur between two cells (top row) or a single cell on glass (bottom row), creating many nearly parallel filaments aligned along the axis of the pull and perpendicular to the cell border. Thick CP pulls (b, n=18) exhibit a wide foot pad at their end which can pull to create many filaments, and nearly 90% exhibit vigorous bubbling (triangular arrows) during the pulling event. Thick CPs are typically aligned with the cell long axis and constitute the entire trailing edge of a cell. Thin CP pulls (c, n=21) are significantly thinner (p<0.05) than thick CP pulls, could be at any angle with respect to the cell long axis, and do not generally constitute the entire trailing edge of a cell. Nearly 50% exhibit minor bubbling, either by the CP itself or by the collaborating cell at the base of the forming filament. Ultrafine CP pulls (d, n=5) are created by the thinnest and longest CPs which are reminiscent of tunneling nanotubes. They always extended between 2 cells, and 80% contained gondola-like vesicles (triangular arrows) that were transported with the pull. Cell surface pulls (e, n=36) occur when the cell rapidly contracts its trailing edge as it migrates across the field of view (bottom row), or when two adjacent, parallel cells rapidly separate (top row), and filaments appear to originate directly from nucleation points on the cell surface. CP and filaments at time=0 are circled and the direction of the pull is indicated by the large arrows. The dashed rectangle shows the measured filament. Scale bars are 10 μm, time is min:sec.

**Supplemental Fig 2:**
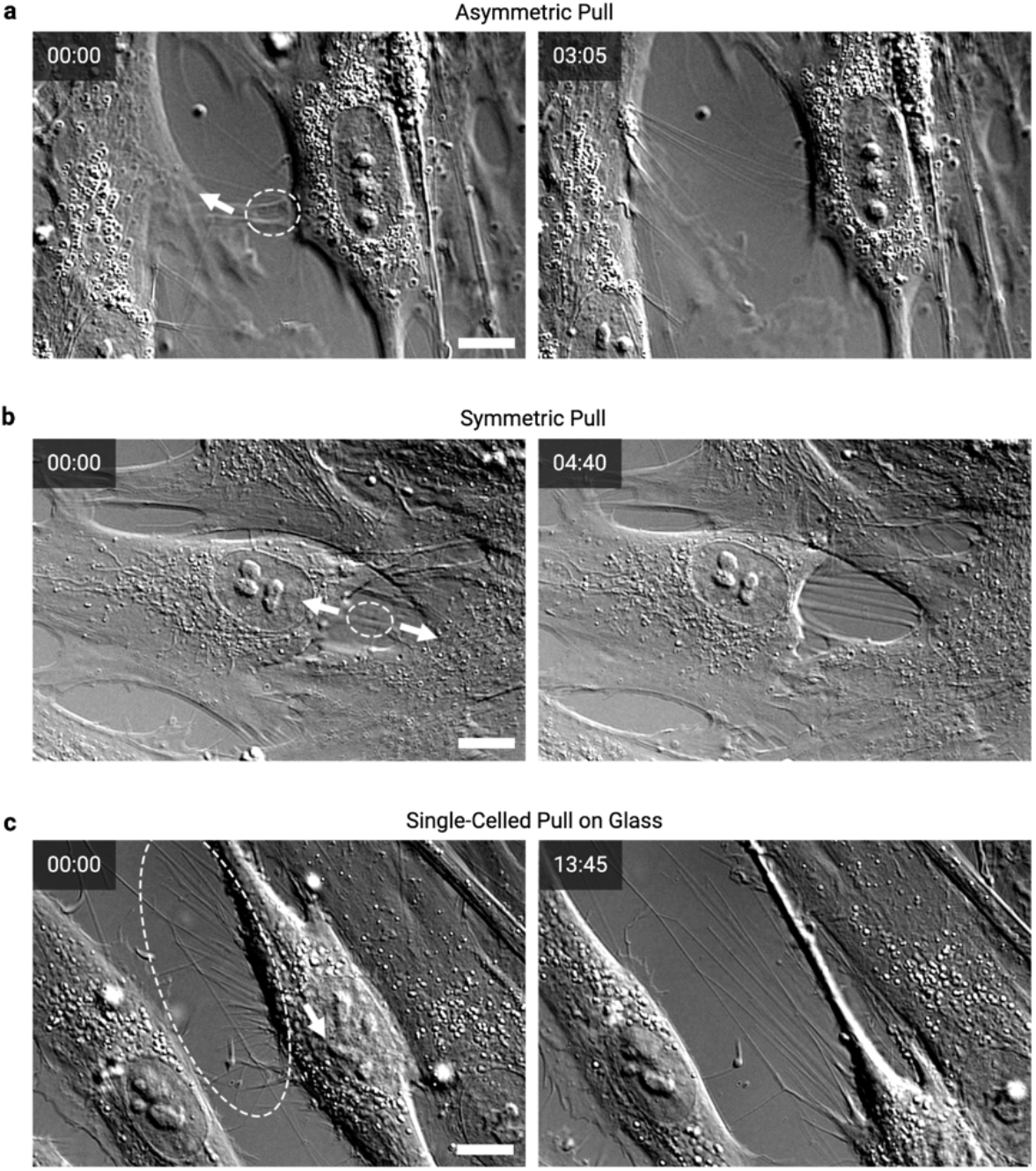
Examples of asymmetric, symmetric, and single-celled pulls. a, Representative asymmetric pull where the left cell process pulls much more quickly than the right cell. b, Representative symmetric pull where the left cell process pulls at the same speed as the cell on the right. c, Representative single-celled pull on filaments that are anchored to the glass. All forming filaments started and ended at fixed points, either on a cell surface, cell process, or the glass. Initial filaments are circled and the arrows indicate the direction of the pulls. Scale bars are 10 μm, time is min:sec.

**Supplemental Fig 3.**
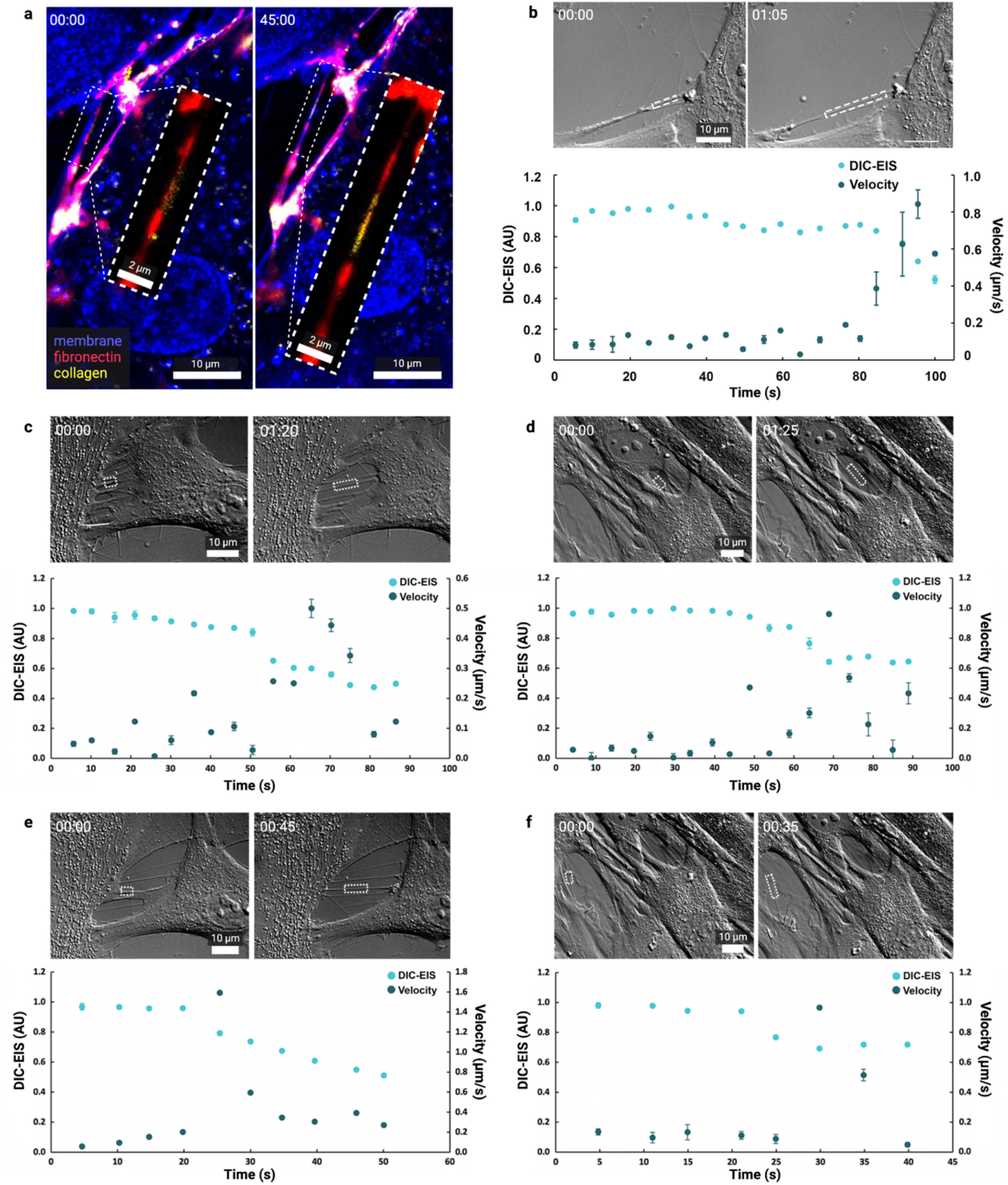
Filaments thin during the pull. a, Live-cell PHCF confocal images of a filament stretching and thinning during a pull. Cells were stained for nucleus (blue), membrane (blue), fibronectin (red), and collagen (yellow). Inset magnifies the central portion of the filament, displaying only fibronectin and collagen. At time=0, the membrane and fibronectin appear thicker and the collagen appears more diffuse. After 45 minutes, the membrane appears thinner, fibronectin appears thinner and stretched, and collagen appears brighter and more concentrated along the filament. b-f, DIC images of a filament stretching and thinning during a pull. Plots show the change in filament thickness (measured using DIC-EIS) and velocity of the pull during the pulling event. The filaments analyzed became thinner during the pulling event, and generally, an increase in pull velocity corresponded to a decrease in thickness. Rectangles indicate the area of the filament that was analyzed, time is min:sec.

**Supplemental Figure 4:**
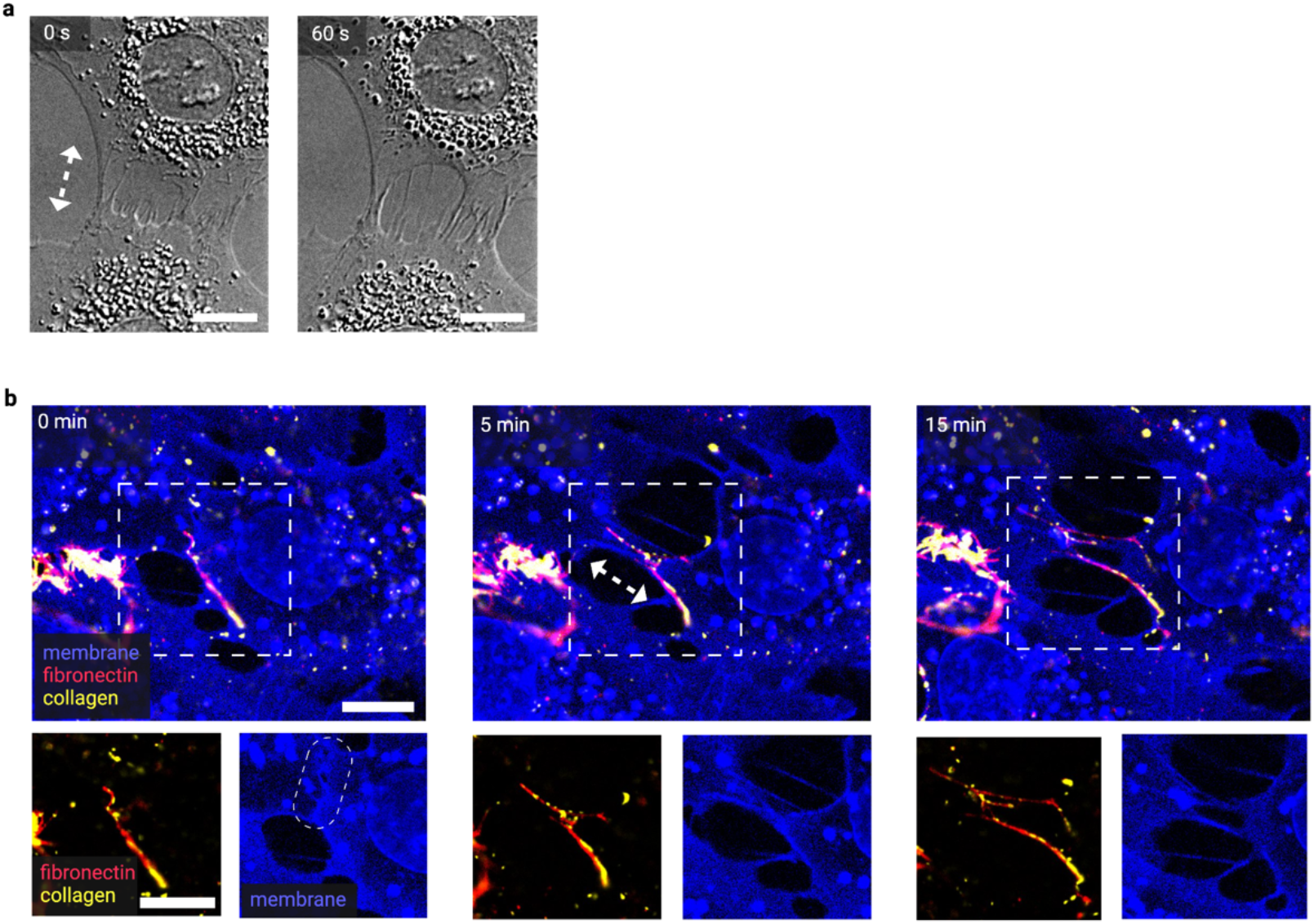
Rat pull. a, DIC live-cell time-lapse imaging of rat fibroblasts. Arrow indicates the direction of the pull. b, Confocal live-cell time-lapse imaging of rat fibroblasts stained for nucleus (blue), membrane (blue), fibronectin (red), and collagen (yellow). Inset magnifies the pull and displays the ECM protein channels separate from the membrane channel. Initially, the two cell surfaces are touching (dashed circle) and fibronectin and collagen are concentrated on the cell surface. As the cells pull (the large arrow indicates pull direction), fibronectin and collagen are stretched into ECM filaments that have a membrane filament at their core. Scale bars are 10 μm.

**Supplemental Figure 5:**
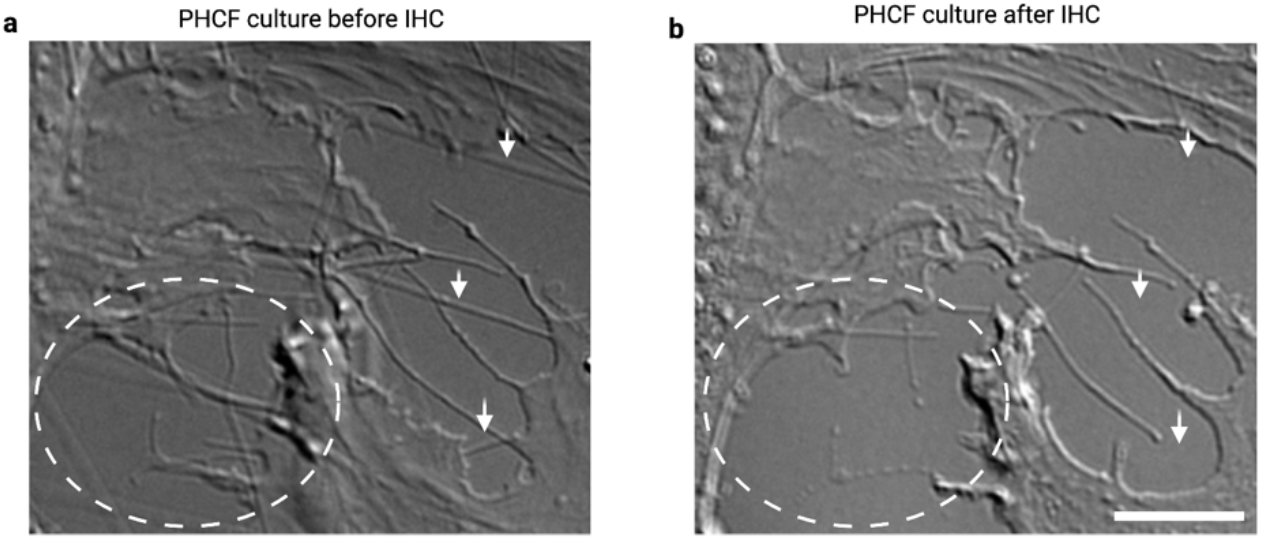
IHC damages filaments. a, Live-cell DIC image of a PHCF culture after a pull, where many filaments are visible (arrows and circles). b, DIC image of PHCF culture after IHC, showing the damage and destruction that occurs to the filaments (arrows and circles). Scale bar is 10 μm.

**Supplemental Figure 6:**
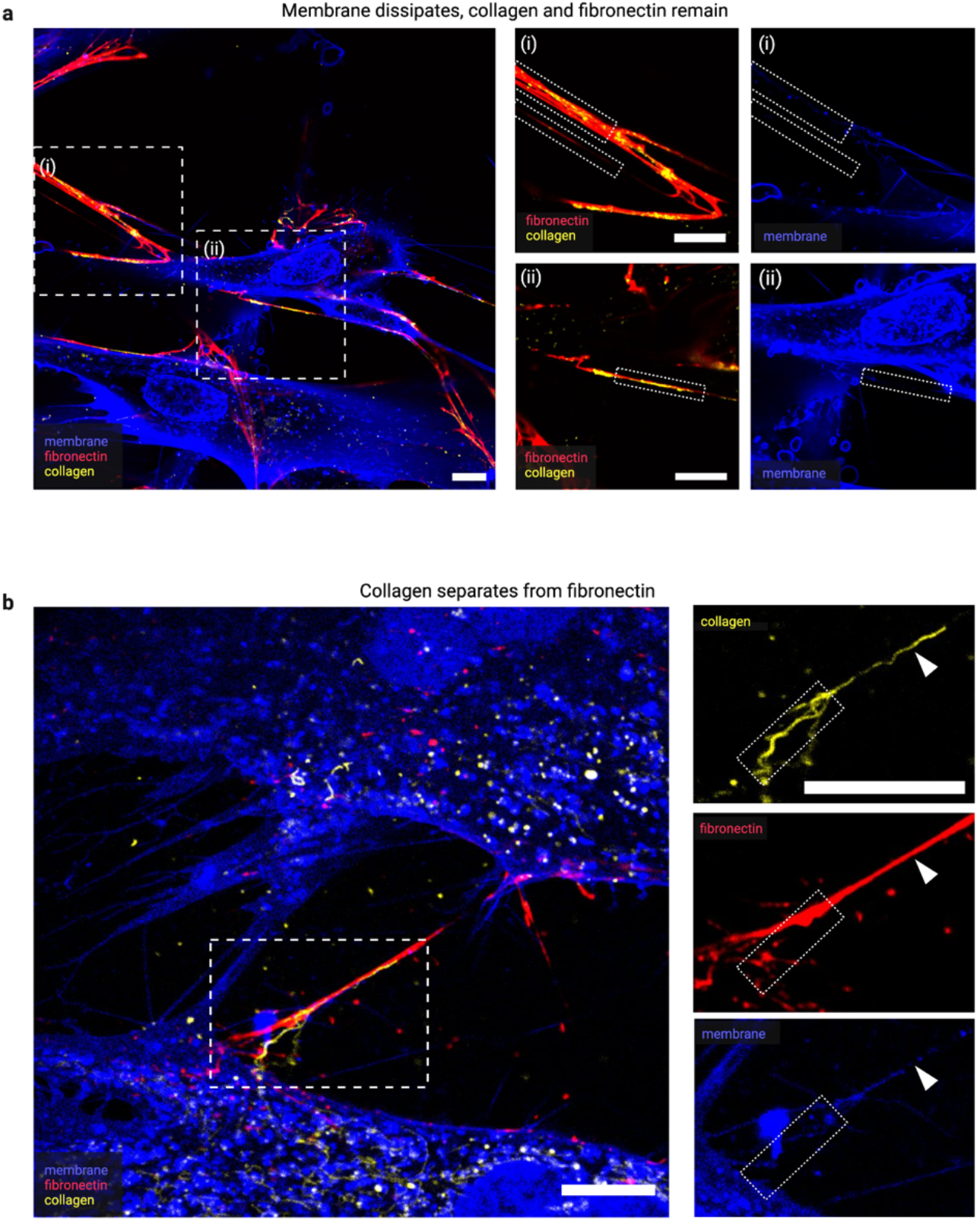
col separates from FN and membrane dissipates. Live-cell confocal images of PHCFs stained for nucleus (blue), membrane (blue), fibronectin (red), and collagen (yellow). a, Low magnification image shows a cell pulling on ECM filaments. i-ii, Prominent filaments comprising fibronectin and punctate collagen (small rectangles) do not contain membrane, suggesting that the membrane dissipates after facilitating filament formation. b, Low magnification image shows a filament stretched between two cells. The main section of the filament (arrow) primarily comprises fibronectin and little membrane. A collagen filament appears bound to fibronectin in the main section (arrow), but appears to be separating from the filament at the base (small rectangle). Scale bars are 10 μm.

## Supplemental Video Legends

**Supplemental Video 1. Videos of five pull types with corresponding velocity and extensional strain rate plots.**

Representative PHCF DIC live-cell time-lapse videos of each pull type and their corresponding kinematics plots of velocity and extensional strain rate. Kinematics data is presented as mean ± standard deviation (n = 3 observers), and the plots are standardized so maximum velocity and extensional strain rate occur at time = 0. Velocities and extensional strain rates were measured by tracking the length of filaments (shown as brackets) during the pulling event.

**Supplemental Video 2: Additional examples of each pull type.**

Representative PHCF DIC live-cell time-lapse videos of additional pull type examples. Flat CP pulls are created by flat, transparent, lamellipodia-like projections on the cell surface. Thick CP pulls are aligned along the long cell axis and exhibit a wide, bubbling foot pad. Thin CP pulls are significantly thinner than thick CP pulls and can exhibit minor bubbling. Ultrafine CP pulls are created by the thinnest and longest CPs, reminiscent of tunneling nanotubes. Cell surface pulls occur when the cell rapidly contracts its trailing edge, or when two adjacent cells rapidly separate, and filaments originate directly from nucleation points on the cell surface. Time is min:sec.

**Supplemental Video 3: Examples of asymmetric, symmetric, and single-celled pulls.**

Representative PHCF DIC live-cell time-lapse videos of additional asymmetric, symmetric, and single-celled pulling examples. In asymmetric pulls, one cell process pulls much more quickly than its partner cell. In symmetric pulls, both cells pull at approximately the same speed. In single-celled pulls, one cell pulls on filaments that are anchored to the glass. Time shown as min:sec.

**Supplemental Video 4: Notable observations that occur during pulling events.**

Representative DIC live-cell time-lapse videos of PHCFs showing examples of notable observations that occur during pulling events. In some pulls, vesicles form immediately before the pulling event and concentrate at the cell surface. As the cell pulls and filaments form, the vesicles appear to deplete. Filaments can also be reeled into the cell. Additionally, cells can pull in two, often nearly orthogonal, directions.

**Supplemental Video 5: Pull causes ECM structure formation.**

Live-cell confocal time-lapse images of a Day 1 PHCF culture stained for nucleus (blue), membrane (blue), fibronectin (red), and collagen (yellow). A slow cell surface pull causes the formation of ECM. First, punctate collagen and fibronectin globs become brighter and more abundant. As the pull begins, fibronectin appears to streak in the direction of the pull. Fibronectin becomes brighter and begins forming a matrix of filaments. Collagen colocalizes with fibronectin and elongates with the stretching ECM. By enhancing the membrane, we can see a network of membranous cell filaments colocalized with most of the newly-formed ECM.

**Supplemental Video 6: Two proposed models for mechanically driven filament assembly.**

Live-cell imaging examples of two proposed models for mechanically driven filament assembly. Confocal time-lapse images are of PHCFs stained for nucleus (blue), membrane (blue), fibronectin (red), and collagen (yellow). Model 1 shows filament formation without a cell membrane extension at the core. Here, a fibronectin filament streaks out of a vesicle (arrow) as the cell bubbles and pulls. The end section of the filament comprises only fibronectin and no membrane. Model 2 shows filament formation with a cell membrane extension at the core. Here, a filament comprising membrane, fibronectin, and collagen is stretched between two adjacent cells.

